# Identification of novel mutations associated with clofazimine resistance in *Mycobacterium abscessus*

**DOI:** 10.1101/276022

**Authors:** Yuanyuan Chen, Jiazhen Chen, Shuo Zhang, Wanliang Shi, Wenhong Zhang, Min Zhu, Ying Zhang

**Author notes:** Corresponding author: Ying Zhang, Department of Molecular Microbiology and Immunology, Bloomberg School of Public Health, Johns Hopkins University, Baltimore, MD 21205, USA. Contributed equally (Yuanyuan Chen, Jiazhen Chen).

## Abstract

*Mycobacterium abscessus* (*Mab*) is a major non-tuberculous mycobacterial (NTM) pathogen responsible for about 80% of all pulmonary infections caused by rapidly growing mycobacteria. Clofazimine is an effective drug active against *Mab* and shows synergistic activity when given with amikacin, but the mechanism of resistance to clofazimine in *Mab* is unknown.

**Objective:** To investigate the molecular basis of clofazimine resistance in *Mab.*

**Methods:** We isolated 29 *Mab* mutants resistant to clofazimine, and subjected them to whole genome sequencing and Sanger sequencing to identify possible mutations associated with clofazimine resistance.

**Results:** Mutations in MAB_2299c gene which encodes possible transcriptional regulatory protein were identified in 23 of the 29 clofazimine-resistant mutants. In addition, 6 mutations in MAB_1483 were found in 21 of the 29 mutants, and one mutation in MAB_0540 was found in 16 of the 29 mutants. Mutations in MAB_0416c, MAB_4099c, MAB_2613, MAB_0409, MAB_1426 were also associated with clofazimine resistance in less frequency. Two identical mutations which are likely to be polymorphisms unrelated to clofazimine resistance were found in MAB_4605c and MAB_4323 in 13 mutants.

**Conclusion:** Mutations in MAB_2299c, MAB_1483, and MAB_0540 are the major mechanisms of clofazimine resistance in *Mab*. Future studies are needed to address the role of the identified mutations in clofazimine resistance in *Mab*, and our findings have implications for developing a rapid molecular test for detecting clofazimine resistance in this organism.

## Introduction

*Mycobacterium abscessus* (*Mab*) complex is a group of rapidly growing nontuberculous mycobacteria (NTM) that can cause severe human diseases, including respiratory, skin and soft tissue disorders, particularly in cystic fibrosis and elderly patients.^[1]^ *Mab* is a major NTM pathogen that is responsible for about 80% of all pulmonary infections caused by rapidly growing NTM. *Mab* complex is notoriously resistant to standard antituberculous agents and most antimicrobial agents.^3^ It is also resistant to disinfectants and, therefore, can cause post-surgical and post-procedural infections.^3,4^ *Mab* has been classified into three subspecies on the basis of *rpoB* sequences: *M*. *abscessus* (sensu stricto), *M*. *massiliense*, and *M*. *bolletii.*^1–3^ They have different suscepbility to antimicrobial agents, and *Mab* subsp. *abscessus* is the most prevalent and resistant and is called “nightmare bacteria”.^3^ The treatment regimen for *Mab* disease is a combination of amikacin, cefocetin, clarithromycin and imipenam for the initial phase and oral antimicrobials for the continuation phase, for a total duration of 12-24 months.^2^ With such a complex regimen and long duration, the outcome of *Mab* disease is still poor. It has been reported that up to 67% of patients’ treatment failed.^5^

Clofazimine, a drug that is currently used for leprosy and multidrug-resistant tuberculosis treatment,^6^ is also active against *Mab* and shows synergistic activity when given together with amikacin.^7–10^ However, its mechanism of action in *Mab* remains to be established. Intracellular redox cycling and membrane destabilization and dysfunction could be the mechanism of clofazimine-mediated antimicrobial activity.^6^ Furthermore, the mechanism of resistance to clofazimine in *Mab* is unknown.

To better understand the mechanism of clofazimine resistance and to develop more rapid molecular tests for detection of its resistance, we characterized 29 clofazimine-resistant mutants isolated in vitro from *M*. *abscessus* ATCC 19977 and found several new genes that are distinct from the known resistance mechanisms in *M. tuberculosis* ^11^ that are associated with clofazimine resistance in *Mab.*

## Materials and methods

### Isolation of *Mab* mutants resistant to clofazimine and clofazimine susceptibility testing

*M. abscessus* ATCC 19977 was grown in 7H9 liquid medium (Difco) supplemented with 0.05% Tween 80 and 10% bovine serum albumin-dextrose-catalase (ADC) enrichment at 37 □ for 3 days (exponential phase), and was diluted 100 times. Clofazimine (Sigma-Aldrich Co.) was dissolved in dimethyl sulfoxide (DMSO, Difco) at a stock concentration of 5 mg/mL and incorporated into 7H11 agar plates containing albumin-dextrose-catalase (ADC) supplement at concentrations of 1, 2, 4 or 8 mg/L. Aliquots of diluted ATCC 19977 (1:100 dilution) was spead on clofazimine containing plates, and also on 7H11 agar plates to confirm the CFU count. Mutants that grew on the plates containing 8 mg/L clofazimine after 5 day incubation at 37 □ were picked and grown in 7H9 liquid medium for confirming clofazimine resistance. The clofazimine susceptibility testing of the clofazimine-resistant mutants was performed on 7H11 agar plates containing 2, 4, 8, 16, 32 mg/L clofazimine. Wild type *M*. *abscessus* ATCC 19977 was included as a drug susceptible control strain for the clofazimine susceptibility testing, and did not grow on plates containing 2, 4, 8, 16, 32 mg/L clofazimine, while the clofazimine-resistant mutants grew on clofazimine containing plates.

### Whole genome sequencing

Genomic DNA was isolated from bacterial cultures (3 ml) as described previously.^12^ Briefly, the bacterial cells were heat killed by incubating them at 80 □ for 20 min followed by glass bead (diameter, 0.1 mm; Sigma) disruption by vigorous vortexing at high speed for 5 min. Bacterial lysates were extracted with phenol-chloroform-isoamyl alcohol (25:24:1). Genomic DNA was precipitated with 2 volumes of absolute alcohol, collected by centrifugation, washed with 70% alcohol, and then air-dried. The genomic DNA samples from 13 clofazimine resistant mutants were subjected to whole genome sequencing using Illumina HiSeq 2000 machine. Paired-end sequencing libraries for genomic DNA of each strain were barcoded and constructed with insert sizes of approximately 300 bp using TruSeq DNA Sample Preparation kits (Illumina, USA) according to manufacturer’s instructions. For each strain, 1.0 G-1.5 G bases (200-fold to 300-fold genome coverage) were generated after barcodes were trimmed. High-quality data were aligned with the reference sequence of *Mab* subsp. *abscessus* ATCC 19977 (NC_010397.1) using SOAPaligner. Only reads where both ends aligned to the reference sequence were used for single nucleotide variant (SNV) and insertion and deletion (InDels) analysis. SNVs and InDels ranging from 1 to 5 bp were sorted and called at minimum reads of 10.

### Polymerase chain reaction (PCR) and DNA sequencing

The genomic DNA from clofazimine-resistant mutants of *Mab* isolated in vitro was subjected to PCR amplification using MAB_4323 primers MAB_4323F (5‘-CTATCCGAGCACACGCTTACC -3‘; 331 bp before InDel) and MAB_4323R (5‘-GCGATGTCAATGAGGAGGAG -3‘; 304 bp after InDel). Qiagen HotStar Taq^R^ DNA Polymerase was applied for PCR amplification and parameters as follows: heat denaturation at 95 □ 15 min followed by 30 cycles of 94 □ 1 min, 51.5 □ 1 min, 72 □ 1 min followed by extension at 72 □ for 10 min. The MAB_4605c gene was PCR-amplified using primers MAB_4605cF (5‘-GACAACACCGTCCGCCATT -3‘; 349 bp before SNV) and MAB_4605cR (5‘-GATCGGCTGGTCGGATACCT -3‘; 363 bp after SNV). The MAB_4605c PCR products were obtained using the parameters of PCR amplification as follows: heat denaturation at 95 □ 15 min followed by 30 cycles of 94 □ 1 min, 53 □ 1 min, 72 □ 1 min followed by extension at 72 □ for 10 min. All the PCR products were purified according to the instructions of QIAquick PCR Purification Kit, then sequenced by Macrogen USA Corp. to confirm the mutations in those genes in selected mutants.

## Results

### MIC determination for *Mab* ATCC 19977

The minimal inhibitory concentration (MIC) of clofazimine for the *Mab* ATCC 19977 strain was determined by the agar diluted method on 7H11 agar plates containing 0.5, 1, 2, 4, 8 mg/L clofazimine. Log phase ATCC 19977 culture grown in 7H9 medium was diluted 1000 times, 10 µl inolula (approximate 2×10^4^ CFU) were added on plates containing different concentrations of clofazimine, and also on 7H11 agar plate as a control. After 3 days of incubation at 37C, the growth of the Mab ATCC 19977 on plates containing 2, 4, 8 mg/L clofazimine was inhibited, indicating that the MIC of clofazimine was 2 mg/L.

### Isolation of *Mab* mutants resistant to clofazimine

Approximately 2×10^7^ of Mab ATCC 19977 was spread on plates containing different concentrations of clofazimine for mutant isolation. After 5-day incubation, 29 mutants were obtained on the plates containing 8 mg/L clofazimine. The mutation frequency of resistant mutants to 8 mg/L clofazimine was about 5×10^-6^. We then determined the MICs for all the mutants on 7H11 agar plates containing 2, 4, 8, 16, 32 mg/L clofazimine with the wild type *Mab* ATCC 19977 as a drug susceptible control strain. We found that all the *Mab* mutants grew on all clofazimine containing plates, indicating that their MICs of clofazimine were greater than 32 mg/L, while the wild type *Mab* grew only on no drug control plate but not on clofazimine containing plates.

### Whole genome sequencing identified novel genes associated with clofazimine resistance in *Mab*

To identify possible new mechanisms of clofazimine resistance, we subjected the 29 clofazimine-resistant mutants to whole genome sequencing. Comparative genome sequence analyses revealed that 23 of 29 CFZ-resistant mutants had mutations in MAB_2299c encoding the transcriptional repressor (Table 1). It is interesting to note that most (18 of 23) of the mutations in MAB_2299c are loss of function mutations due to InDels or stop codons with only 5 SNVs that caused amino acid changes. We also found 6 mutations in MAB_1483 gene in respective 20 mutants (N1-N14, N16, M1, M11, M19, M29 and M84) including 5 InDels causing frameshift mutations, 1 SNV that results in stop codons (Table 1). The other mutant, M65, had a C554T mutation, causing the amino acid change A185T in the MAB_0416c gene, which encodes putative Crp/Fnr-family transriptional regulator (Table 1). In addition, two other mutants with respective mutations in MAB_4099c encoding probable non-ribosomal peptide synthetase and MAB_2613 encoding putative glucose/mannose:H+ symporter GlcP were identified, one having a T9776C mutation causing Y3259C amino acid change and one having a T310C mutation causing V104A amino acid change (Table 1).

**Table 1.** Mutations identified from 29 clofazimine resistant *M. abscessus* mutants by whole genome sequencing

| Gene | Nucleotide mutation | Amino acid change | Mutation type | Gene_product | Strains No. |
| --- | --- | --- | --- | --- | --- |
| MAB_2299c | Ins GGC (132) | Ins 44 | Insertion | Possible transcriptional regulatory protein | N1, N5, N6, N12 |
| MAB_2299c | Ins G (276) | FSC92 | Insertion | Possible transcriptional regulatory protein | N2 |
| MAB_2299c | Del G (322) | FSC108 | Deletion | Possible transcriptional regulatory protein | N3, N8 |
| MAB_2299c | Ins TT (344) | FSC115 | Insertion | Possible transcriptional regulatory protein | N4, N7 |
| MAB_2299c | Ins G (223) | FSC75 | Insertion | Possible transcriptional regulatory protein | N10 |
| MAB_2299c | Del C (215) | FSC72 | Deletion | Possible transcriptional regulatory protein | N16 |
| MAB_2299c | C63A | E21STOP | Deletion | Possible transcriptional regulatory protein | M19 |
| MAB_2299c | T518C | H173R | SNV | Possible transcriptional regulatory protein | M1, M2, M7 |
| MAB_2299c | C642A | A214S | SNV | Possible transcriptional regulatory protein | M8 |
| MAB_2299c | Ins G (391) | FSC131 | Deletion | Possible transcriptional regulatory protein | M3 |
| MAB_2299c | Ins G (345) | FSC115 | Insertion | Possible transcriptional regulatory protein | M11, M23, M29 |
| MAB_2299c | Ins A (494) | FSC165 | Insertion | Possible transcriptional regulatory protein | M14 |
| MAB_2299c | G19A | Q7STOP | Deletion | Possible transcriptional regulatory protein | N11 |
| MAB_2299c | C329T | C110Y | SNV | Possible transcriptional regulatory protein | N14 |
| MAB_1483 | Del G (708) | FSC236 | Deletion | hypothetical protein | N1-N16 |
| MAB_1483 | G363A | W121STOP | Deletion | hypothetical protein | M1 |
| MAB_1483 | Del TACCG (489) | FSC163 | Deletion | hypothetical protein | M84 |
| MAB_1483 | Ins G (316) | FSC106 | Insertion | hypothetical protein | M29 |
| MAB_1483 | Ins T (104) | FSC35 | Insertion | hypothetical protein | M11 |
| MAB_1483 | Del A (343) | FSC115 | Deletion | hypothetical protein | M19 |
| GlpK | Ins GGC (3'+28bp) | - | 3' intergenic insertion | Glycerol kinase | N11 |
| MAB_0540 | Del G (-45) | - | 5' intergenic deletion | hypothetical protein | N1-N16 |
| MAB_0416c | C554T | A185T | SNV | Putative Crp/Fnr-family transcriptional regulator | M65 |
| MAB_4099c | T9776C | Y3259C | SNV | Probable non-ribosomal peptide synthetase | M11 |
| MAB_2613 | T310C | V104A | SNV | Putative glucose/mannose:H<sup>+</sup> symporter GlcP | M7 |
| MAB_0409 | T(-88)G | - | 5' SNV | Putative transcriptional regulator WhiB | N13 |
| MAB_1426 | A649C | K217Q | SNV | Putative cytochrome P450 | N3, N10 |
| MAB_4605c | G122A | A41V | SNV | Probable aldehyde dehydrogenase | M1-M3, M7, M8, M11, M14, M19, M23, M29, M34, M65, M84 |
| MAB_4323 | Ins C (1493) | FSC498 | Insertion | Putative amino acid transporter | M1-M3, M7, M8, M11, M14, M19, M23, M29, M34, M65, M84 |

Whole genome sequencing indicated that G122A mutation in MAB_4605c encoding probable aldehyde dehydrogenase and C1493 insertion in MAB_4323 encoding amino acid transporter were the most common mutations in 13 clofazimine resistant mutants (Table 1). To confirm these two mutations to be genuine, PCR using MAB_4605c and MAB_4323 primers was performed on six individual mutants (M11, M23, M29, M34, M65, M84) and clofazimine susceptible control strain (ATCC 19977), and Sanger sequencing of the PCR products indicated that the two mutations in the 6 mutants were correct.

## Discussion

Although clofazimine was discovered in 1957 as a riminophenazine drug for treating TB,^[12]^ it is mainly used for the treatment of leprosy ^14^ and also as a core second-line agent for the treatment of MDR-TB.^15^ In *M. tuberculosis*, the mechanisms of resistance to clofazimine are mainly due to mutations in the transcriptional repressor Rv0678, with concomitant upregulation of the efflux pump, MmpL5, and occassionally in Rv1979c (a permease) or Rv2535c (PepQ).^11,16^ However, the mechanisms of action and resistance of clofazimine in *Mab* has not been reported so far. In this study, we isolated *Mab* mutants resistant to clofazimine and found several genes (MAB_2299c, MAB_1483, MAB_0540, MAB_0416c, MAB_4099c, MAB_2613, MAB_0409, MAB_1426) whose mutations are associated with clofazimine resistance through comparative genome sequence analyses.

It is worth noting that 23 of 29 mutants had InDels or SNVs in the MAB_2299c gene encoding possible transcriptional regulatory protein. MAB_2299c is a homolog of *M. tuberculosis* Rv0452, a putative AcrR family transcription repressor of efflux genes *mmpL4* and *mmpS4* (Fig. 1). Rv0452 and *mmpL4-mmpS4* are organized in the same genomic structure as Rv0678 and *mmpL5-mmpS5* in *M. tuberculosis* (see Fig. 1). Therefore, mutations in MAB_2299c in *Mab* would be analogous to mutations in Rv0678 in *M. tuberculosis* in causing clofazimine resistance. Future studies are required to confirm this.

**Fig. 1.**
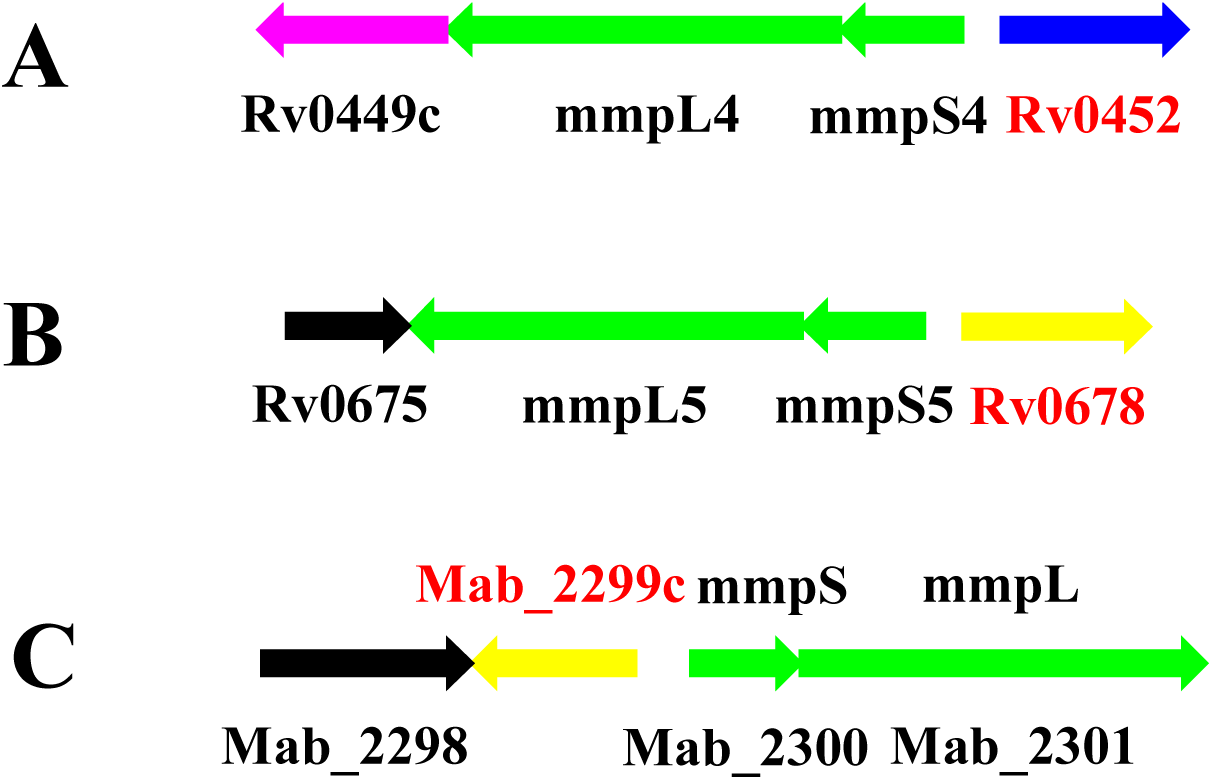
Similarity of genomic organization structures of Mab_2299c and Rv0452 and Rv0678. Mab_2299c is the homolog of Rv0452, which is a transcriptional repressor of the transmembrane MmpS4-MmpL4 efflux pump in *M. tuberculosis* (A). Rv0452 has similar genomic organization as Rv0678 which is known to be a transcription repressor of MmpS5-MmpL5 efflux pump (B) involved in clofazimine resistance in *M. tuberculosis.* The similar genomic structure of Mab_2299c and its adjacent mmpS and mmpL genes in *Mab* (C) to Rv0452 and Rv0678 suggests that Mab_2299c mutations have the same mechanism of CFZ resistance as that in Rv0678 mutation in *M. tuberculosis*.

Twenty-one of 29 mutants had loss of function mutaions in MAB_1483 encoding a hypothetical protein with no significant homologs in *M. tuberculosis*. Sixteen of 29 mutants had a -45 G deletion in the upstream of MAB_0540, but its function is unknown. The role and function of MAB_1483 and MAB_0540 need to be explored with regard to clofazimine resistance. MAB_0416c encodes a putative Crp/Fnr-family transcriptional regulator, and mutation in its homolog Rv3676 caused growth defect and is involved virulence attenuation in *M. tuberculosis.*^17^ MAB_0409 encodes putative transcriptional regulator WhiB, and its promoter region -88 mutation may affect the transcription of the WhiB homolog and lead to clofazimine resistance, though the mechanisms involved remains to be identified. MAB_4099c encoding probable non-ribosomal peptide synthetase and MAB_2613 encoding putative glucose/mannose:H+ symporter GlcP are less likely causal in clofazimine resistant *Mab*. Although we found G122A mutation in MAB_4605c encoding probable aldehyde dehydrogenase and C1493 insertion in MAB_4323 encoding amino acid transporter in all 13 clofazimine resistant mutants (Table 1), we believe these SNVs may not have a direct role in causing clofazimine resistance as they are present in only one batch of clofazimine resistant mutants (the M series) but not in the second batch of mutants (the N series).

Despite numerous studies, the mechanism of action of clofazimine has remained elusive. The drug is extremely hydrophobic, suggesting that its mode of action is closely associated with its effect on membranes.^6^ Clofazimine reduction by mycobacteria was noted in the first study^13^ on the drug which is supposed to react with molecular O_2_ to form reactive oxygen species (ROS), raising speculation that clofazimine antimycobacterial activity was related to the intracellular redox activity of the dye.^6^ Yano et al. proposed that clofazimine competes with the NDH-2 substrate, menaquinone, which accepts electrons/H donated by NADH and delivered them to electron transport chain (ETC) which in turn uses the electrons/H to reduce O_2_, and that NDH-2 catalyzes clofazimine to a reduced form oxidized by molecular oxygen forming superoxide and H_2_O_2_.^18^ However, in this study, the mutations we identified in *Mab* (Table 1) do not map to redox or ETC, which is similar to the situation in *M. tuberculosis.* Thus, the mutation approach may not always be useful for identifying drug targets needed for understanding the mode of action of the drug.

In summary, we identified several new genes whose mutations are associated with clofazimine resistance in *Mab*. Mutations in MAB_2299c, MAB_1483, and MAB_0540 seem to be major mechanisms of clofazimine resistance. Mutations in MAB_2299c encoding the transcriptional repressor of efflux pump is analogous to that of Rv0678 mutations in causing clofazimine resistance in *M. tuberculosis.* Other new genes involved in clofazimine resistance in *Mab* seem to affect transcription or transport functions. Our findings offer new insights about the mechanisms of resistance to clofazimine and its mode of action in *Mab*. Identification of MAB_2299c, MAB_1483, and MAB_0540 mutations in most clofazimine resistant mutants has implications for rapid molecular detection of clofazimine resistant *Mab*. Future studies are needed to assess the role of the identified mutations as new mechanisms of clofazimine resistance in *Mab*.

## References

[1] Griffith DE. *Mycobacterium abscessus* subsp abscessus lung disease: ’trouble ahead, trouble behind…’. F1000Prime Rep. 2014 Nov 4; 6:107. doi: 10.12703/P6-107.

[2] Griffith DE, Aksamit T, Brown–Elliot BA et al. An official ATS/IDSA statement: diagnosis, treatment, and prevention of nontuberculous mycobacterial diseases. Am. J. Respir. Crit. Care Med. 175, 367–416 (2007).

[3] Nessar R, Cambau E, Reyrat JM, Murray A, Gicquel B. Mycobacterium abscessus: a new antibiotic nightmare. J Antimicrob Chemother. 2012 Apr;67(4):810–8. doi: 10.1093/jac/dkr578. Epub 2012 Jan 30.

[4] Lee MR, Sheng WH, Hung CC, Yu CJ, Lee LN, Hsueh PR. *Mycobacterium abscessus* Complex Infections in Humans. Emerg Infect Dis. 2015 Sep;21(9):1638–46. doi: 10.3201/2109.141634.

[5] Jarand J, Levin A, Zhang L, Huitt G, Mitchell JD, Daley CL. Clinical and microbiologic outcomes in patients receiving treatment for *Mycobacterium abscessus* pulmonary disease. Clin Infect Dis. 2011 Mar 1;52(5):565–71. doi: 10.1093/cid/ciq237.

[6] Cholo MC, Steel HC, Fourie PB, Germishuizen WA, Anderson R. Clofazimine: current status and future prospects. J Antimicrob Chemother 2012; 67: 290–298.

[7] Singh S, Bouzinbi N, Chaturvedi V, Godreuil S, Kremer L. In vitro evaluation of a new drug combination against clinical isolates belonging to the *Mycobacterium abscessus* complex. Clin Microbiol Infect. 2014 Dec;20(12):O1124–7. doi: 10.1111/1469-0691.12780. Epub 2014 Oct 3.

[8] van Ingen J, Totten SE, Helstrom NK, Heifets LB, Boeree MJ, Daley CL. In vitro synergy between clofazimine and amikacin in treatment of nontuberculous mycobacterial disease. Antimicrob Agents Chemother 2012; 56: 6324–6327.

[9] Shen GH, Wu BD, Hu ST, Lin CF, Wu KM, Chen JH. High efficacy of clofazimine and its synergistic effect with amikacin against rapidly growing mycobacteria. Int J Antimicrob Agents 2010; 35: 400–404.

[10] Ferro BE, Meletiadis J, Wattenberg M, de Jong A, van Soolingen D, Mouton JW, van Ingen J. Clofazimine Prevents the Regrowth of *Mycobacterium abscessus* and *Mycobacterium avium* Type Strains Exposed to Amikacin and Clarithromycin. Antimicrob Agents Chemother. 2015 Dec 7;60(2):1097–105. doi: 10.1128/AAC.02615- 15. Print 2016 Feb. PubMed PMID: 26643335;

[11] Zhang S, Chen J, Cui P, Shi W, Zhang W, Zhang Y. Identification of novel mutations associated with clofazimine resistance in *Mycobacterium tuberculosis*. J Antimicrob Chemother. 2015 Sep;70(9):2507–10. doi: 10.1093/jac/dkv150. Epub 2015 Jun 4.

[12] Zhang Y, Garcia MJ, Lathigra R, Allen B, Moreno C, van Embden JD, Young D. Alterations in the superoxide dismutase gene of an isoniazid-resistant strain of *Mycobacterium tuberculosis.* Infect Immun. 1992 Jun;60(6):2160–5.

[13] Barry VC, Belton JG, Conalty ML et al. A new series of phenazine (rimino-compounds) with high anti-tuberculosis activity. Nature 1957; 179: 1013–5.

[14] Browne SG, Hogerzeil LM. B663 in the treatment of leprosy. Preliminary report of a pilot trial. Lepr Rev 1962; 33: 6–10.

[15] World Health Organizatin. WHO treatment guidelines for drug-resistant tuberculosis (2016 update). http://www.who.int/tb/areas-of-work/drug-resistant-tb/treatment/resources/en/

[16] Hartkoorn RC, Uplekar S, Cole ST. Cross-resistance between clofazimine and bedaquiline through upregulation of MmpL5 in *Mycobacterium tuberculosis*. Antimicrob Agents Chemother 2014; 58: 2979–81.

[17] Akif M, Akhter Y, Hasnain SE, Mande SC. Crystallization and preliminary X-ray crystallographic studies of *Mycobacterium tuberculosis* CRP/FNR family transcription regulator. Acta Crystallogr Sect F Struct Biol Cryst Commun. 2006 Sep 1;62(Pt 9):873–5. Epub 2006 Aug 11.

[18] Yano T, Kassovska-Bratinova S, Teh JS, et al. Reduction of clofazimine by mycobacterial type 2 NADH: quinone oxido-reductase: a pathway for the generation of bactericidal levels of reactive oxygen species. J Biol Chem. 2011; 286:10276–10287.

